# Anti-CD47 immunotherapy as a therapeutic strategy for the treatment of breast cancer brain metastasis

**DOI:** 10.1101/2023.07.25.550566

**Authors:** Jessica D. Mackert, Elizabeth R. Stirling, Adam S. Wilson, Brian Westwood, Dawen Zhao, Hui-Wen Lo, Linda Metheny-Barlow, Katherine L. Cook, Glenn J. Lesser, David R. Soto-Pantoja

## Abstract

The presence of cell surface protein CD47 allows cancer cells to evade innate and adaptive immune surveillance resulting in metastatic spread. CD47 binds to and activates SIRPα on the surface of myeloid cells, inhibiting their phagocytic activity. On the other hand, CD47 binds the matricellular protein Thrombospondin-1, limiting T-cell activation. Thus, blocking CD47 is a potential therapeutic strategy for preventing brain metastasis. To test this hypothesis, breast cancer patient biopsies were stained with antibodies against CD47 to determine differences in protein expression. An anti-CD47 antibody was used in a syngeneic orthotopic triple-negative breast cancer model, and CD47 null mice were used in a breast cancer brain metastasis model by intracardiac injection of the E0771-Br-Luc cell line. Immunohistochemical staining of patient biopsies revealed an 89% increase in CD47 expression in metastatic brain tumors compared to normal adjacent tissue (p ≤ 0.05). Anti-CD47 treatment in mice bearing brain metastatic 4T1br3 orthotopic tumors reduced tumor volume and tumor weight by over 50% compared to control mice (p ≤ 0.05) and increased IBA1 macrophage/microglia marker 5-fold in tumors compared to control (p ≤ 0.05). Additionally, CD47 blockade increased the M1/M2 macrophage ratio in tumors 2.5-fold (p ≤ 0.05). CD47 null mice had an 89% decrease in metastatic brain burden (p ≤ 0.05) compared to control mice in a brain metastasis model. Additionally, RNA sequencing revealed several uniquely expressed genes and significantly enriched genes related to tissue development, cell death, and cell migration tumors treated with anti-CD47 antibodies. Thus, demonstrating that CD47 blockade affects cancer cell and tumor microenvironment signaling to limit metastatic spread and may be an effective therapeutic for triple-negative breast cancer brain metastasis.

## Introduction

Triple-negative breast cancer (TNBC) accounts for approximately 15% of breast cancer cases and is characterized by the absence of estrogen receptor (ER), progesterone receptor (PR), and human epidermal growth factor 2 receptor (HER2) expression (1). Among all the breast cancer subtypes, TNBC has the highest rate of distant metastasis and is associated with the shortest overall survival (OS)(2). Despite undergoing surgical resection and adjuvant chemotherapy, half of all primary TNBC patients in which the tumor is confined to the breast and lymph nodes have a distant recurrence within five years and are prone to metastasis to the central nervous system (CNS) and internal organs(3). Additionally, TNBC patients are more likely to develop brain metastases than other breast cancer subtypes. Brain metastasis (BM) significantly increases patient mortality rate compared with metastasis to other organs, which can be as high as 80% within one year(4). Advances in treatment for patients with ER+/PR+ and/or HER2+ breast cancers have resulted in vast improvements in survival(5); however, TNBC lacks specific targets, and the development of targeted therapies for patients has been lacking.

The recent clinical development of immune checkpoint inhibitors (ICIs) has dramatically changed the landscape of cancer treatment for TNBC(6,7). ICIs targeting inhibitory receptors can induce complete and long-lasting tumor immunity in patients with metastatic and treatment-refractory cancers. Despite the promise of ICIs for TNBC, only a fraction of patients benefit from treatment. Cancer is highly heterogeneous and exploits various mechanisms to evade immune surveillance (8). CD47, also known as integrin-associated protein, is a transmembrane protein that is highly expressed on the surface of various cancer cells. It interacts with thrombospondin-1, signal-regulatory protein-α (SIRP-α), and integrins to regulate multiple cellular functions. In many malignancies, its expression correlates with an aggressive phenotype and an overall poor clinical prognosis (9). Inhibition of CD47 signaling enhances macrophage phagocytic activity and, in preclinical models, leads to impaired tumor growth, inhibition of metastatic spread, and tumor regression.

Several CD47 antagonists have been developed and show great promise in pre-clinical studies. Blockade of the CD47–SIRPα interaction reprograms the tumor microenvironment toward a pro-inflammatory phenotype, which increases cancer cell phagocytosis, antigen presentation, and subsequent T cell activation (10,11). CD47 antagonists are currently in clinical trials and show promising results in both solid tumors and hematological malignancies (12-15). However, little attention has been given to CD47 blockade to treat or prevent metastasis.

In this report, we showed that blockade of CD47 differentially regulates gene expression associated with metastatic spread. Our in vivo data show that the blockade of CD47 may directly target mechanisms that mediate cell migration or that depletion of this receptor in the tumor microenvironment may clear or limit the spread of metastatic cells. Therefore, targeting CD47 should be considered as a potential strategy to limit the brain metastatic burden associated with invasive breast cancer.

## Methods

### Immunohistochemistry

Tissues were fixed in 4% paraformaldehyde for 24 h and immersed in 70% ethanol before paraffin embedding and sectioning at 5 µm. Patients punch biopsies were obtained in a tissue array (Biomax, Cat# GL861) and stained with anti-CD47 antibody B6H12 (Thermo Fischer, Cat#14-0479-82). Tumor tissue sections were stained for IBA1. Briefly, slides were deparaffinized in xylene, rehydrated through graded alcohols, and rinsed in water. Antigen retrieval was performed using citrate buffer (Biogenex, Cat# HK086-9K). Slides were blocked with Dako Serum-Free Protein block (Agilent; Cat # X0909) for 15 min and incubated overnight at 4°C with the primary antibody Ki67 (BD Biosciences, Cat# 558619) and followed staining protocol with DAB as done previously(16). For immunofluorescence the following day, the slides were washed in PBS, counterstained with DAPI (Agilent, Cat# S0809), and mounted with a coverslip. Negative controls with only the secondary antibody were included to account for nonspecific binding. Immunofluorescence was visualized using the Mantra (insert info), and intensity was determined using inForm software (Perkin-Elmer).

### Cell Culture

4T1 and SIMA9 cells were obtained from ATCC (4T1 Cat# CRL-2539; SIM-A9: CRL-3265). 4T1Br-3 was a gift from Dr. Dawen Zhao, and 4T1-Br3-Luc was generated by Dr. Hui-Wen Lo. 4T1 cell lines were cultured in RPMI supplemented with 10% FBS penicillin/streptomycin and glutamine, and SIMA9 cells were cultured in DMEM: F-12 supplemented with heat-inactivated horse serum (5%) and heat-inactivated fetal bovine serum (10%) kept at 37°C and 5% CO2. EO771-Br4-Luc were a gift from Dr. Linda Metheny-Barlow and were maintained in DMEM with 10% FBS, 20 mM HEPES penicillin/streptomycin, and glutamine. IDEXX BioAnalytics performed genetic validation of cell lines.

### *In vivo* models

Orthotopic model: Female Balb/c mice were injected with 4T1-Br3 cells (1 ×10^5^ cells) into the mammary fat pad to induce tumor growth using a 26-gauge needle. Once tumors reached an average of 100 mm^3^, mice were randomized and received IP injections of isotype control (Bioxcell, Cat# BE0089) or anti-CD47 antibody (Bioxcel, Cat# BE0270) every other day for 1 week. Tumor size was measured every third day using a caliper, and the wet weight of the tumors was determined at the end of the study.

Intracardiac model of brain metastasis: Female Balb/c mice (8-10 week old female) were injected with 4T1-Br3-Luc (2x10^5^) by intracardiac injection. After one week, mice were randomized into groups and received IP injections of isotype control or anti-CD47 antibody every other day for 1 week. EO771-Br4 cells were implanted by intracardiac injection (2x10^5^) into C57BL/6 mice or CD47 null mice (8–10-week-old female) in the same background. Brain metastases were determined by IVIS (Perkin Elmer) imaging.

### Flow cytometry

4T1-Par and 4T1-Br3 cells were counted and stained with Fixable Viability Stain 575V (BV605) for 15 minutes at room temperature and washed with PBS+1% FBS. Cells were treated with mouse FcBlock for 10 minutes to limit nonspecific antibody binding, followed by washing. Cells were then stained with an anti-mouse CD47 antibody (Biolegend, Cat# 127504). Cells were acquired using the Fortessa X20 (BD Biosciences), and data were analyzed using FCS Express (De Novo Software).

### Real-Time Measurement of Macrophage Mediated Cytotoxicity

4T1-Br3 breast cancer target cells were seeded into 16-well plates. Cell growth was dynamically monitored using the xCelligence system (Agilent) for 24 h, as we have done previously (17,18). SIMA9 microglia cells were activated for 24h at 37°C with 10 ng/ml of LPS in a complete medium and used at an effector/target ration ratio of 5:1. After addition of effector cells, the analyzer automatically collected measurements. Cell-mediated cytolysis was calculated using the xCELLigence software set to collect impendence data (reported as normalized cell index).

### RNA sequencing

Total RNA was isolated from excised 4T1-Br3 tumor from our in vivo orthotopic model using TRizol reagent protocol (Thermo Fisher, Cat# 15596026). Preparation of library and transcriptome sequencing were performed Novogene Corporation. Gene hits were found using Swiss-Prot, and functional annotation was performed in DAVID and KEGG(19,20). Enrichment analysis for the GO biological process was performed using ShinyGO 0.77(21). Principle component analysis was performed by discrete correlation summation (DCS) as we have done previously(22) to determine the separation of groups by gene expression.

### Statistics

Differences between groups were analyzed by analysis of variance (ANOVA) followed by Fischer’s LSD tests. In vivo, studies were analyzed using repeated-measures ANOVA with SD. An unpaired t-test was used to analyze the differences between two groups. Results are represented as the mean±SD and are considered significant if *p<0.05. Statistical analysis was performed on GraphPad Prism.

## Results

### CD47 increases in brain-trophic triple-negative breast cancer cells and patient metastatic lesions

To determine CD47 expression, flow cytometry was performed on murine 4T1 and 4T1-Br3 TNBC cells (Figure 1A). CD47 surface protein expression was increased 5-fold in 4T1-Br3 brain-trophic cells, compared to parental 4T1 cells (1.000 ± 0.000 vs. 5.304 ± 1.490; Fold-change in 4T1 vs. 4T1-Br3; p=0.045; n=3; Figure 1B). Additionally, immunological staining of paraffin-fixed patient samples revealed that CD47 is increased 1.89-fold in breast cancer brain metastases compared to normal adjacent tissue (34.83 ± 8.669 vs. 65.83 ± 9.912; CD47+ cells in Normal adjacent vs. Brain metastasis; p=0.041; n=4-9; Figures 1C & 1D). We have previously demonstrated that targeting CD47 in the 4T1 cell line reduces TNBC tumor growth in a syngeneic orthotopic model (23). Since we observed upregulation of CD47 over the parental 4T1, we tested whether targeting CD47 would reduce tumor burden in an in vivo orthotopic model using the 4T1Br3 brain trophic cell line. Therefore, female Balb/c mice were injected with 5 x 10^6^ 4T1-Br3 cells in the mammary fat pad. Once tumors reached 100mm^3^, mice were treated with anti-mouse CD47(301) antibody by IP injection every other day for one week. Animal study schematic is shown in Figure 1E. Anti-CD47 treatment significantly reduced tumor volume by 48.4% by day 25 (990.7 ± 87.55 vs. 511.5 ± 38.86; Tumor volume (mm^3^) of IgG vs. CD47(301); p=0.0005; n=6, Figure 1F). Tumor weight was reduced by 41.7% with CD47(301) treatment (0.979 ± 0.074 vs. 0.571 ± 0.059; Tumor weight (g) of IgG vs. CD47(301); p=0.0015; n=6). Therefore, our data shows that blockade of CD47 inhibits brain trophic TNBC orthotopic tumor growth in vivo.

**Figure 1.**
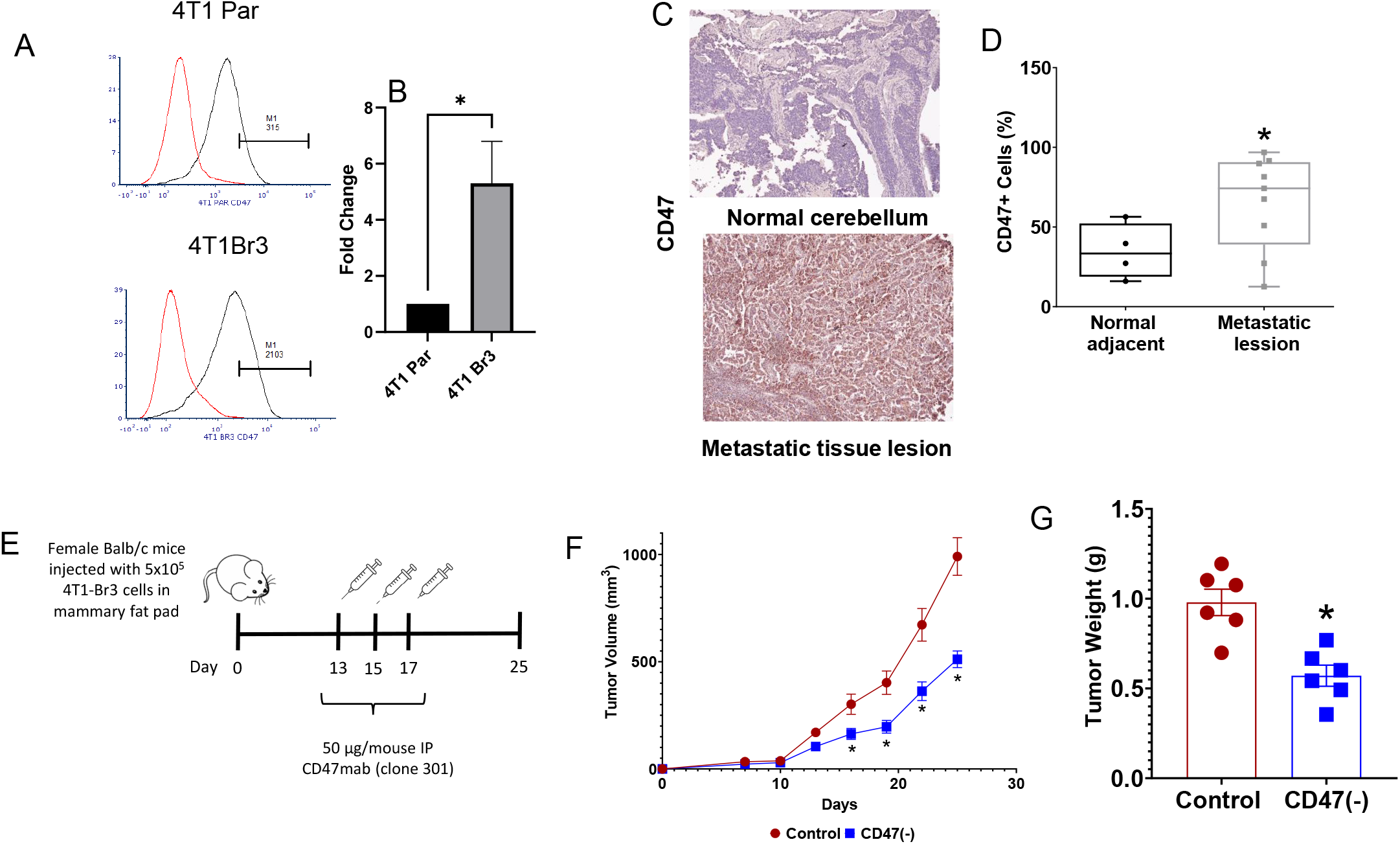
CD47 protein levels in brain metastatic breast cancer cells and patient metastatic lesions. (A) CD47 expression in 4T1-parental vs. 4T1-Br3 (brain trophic) cell lines determined by flow cytometry and fold change increased based on mean fluorescence intensity (n=3, *p<0.05). (C) Patient punch biopsies from normal cerebellum vs. brain metastatic lesions were stained with CD47 anti-human antibody B6H12; (D) sections were scored using Mantra software tissue analysis under bright field (BF) microscopy (n=4-9, *p<0.05). (E) Schematic of in vivo orthotopic 4T1-Br3 model in Balb/c mice treated with isotype (control) or anti-CD47 antibody (CD47(-). (F) Tumor diameter was measured by calipers and tumor volume was calculated using the formula LW^2^/2, and (G) tumors were excised and weighted at the end of the study (n=6, *p<0.06).

### RNA sequencing reveals pathways modulated by CD47 in the tumor microenvironment

At the end of the study, tumors were excised, and a portion of the sample was subjected to analysis by RNA sequencing to understand potential pathways mediating the anti-tumor effect of CD47 blockade. A co-expression Venn diagram shows that over 300 and 250 genes were uniquely detected in anti-CD47 treated and control tumors, respectively (Figure 2A). Furthermore, over 12 500 genes were co-expressed by both groups. Of these genes, 217 were significantly regulated (*p<0.05) between groups, with 116 genes significantly downregulated and 101 upregulated by blockade of CD47 (Figure 2B). Principal component analysis revealed that tumors from mice treated with isotype control or CD47 antibody separated by the differential regulation of gene expression (Figure 3C). Enrichment analysis using the GO Biological process of significantly upregulated genes showed several pathways modulated by anti-CD47 treatment. These include several processes relevant to the metastasis process, including significant enrichment (FDR cutoff 0.05) in pathways related to cell death, motility, and migration. Thus, suggesting potential mechanisms for the observed reduction in in vivo tumor growth observed with CD47 blockade.

**Figure 2.**
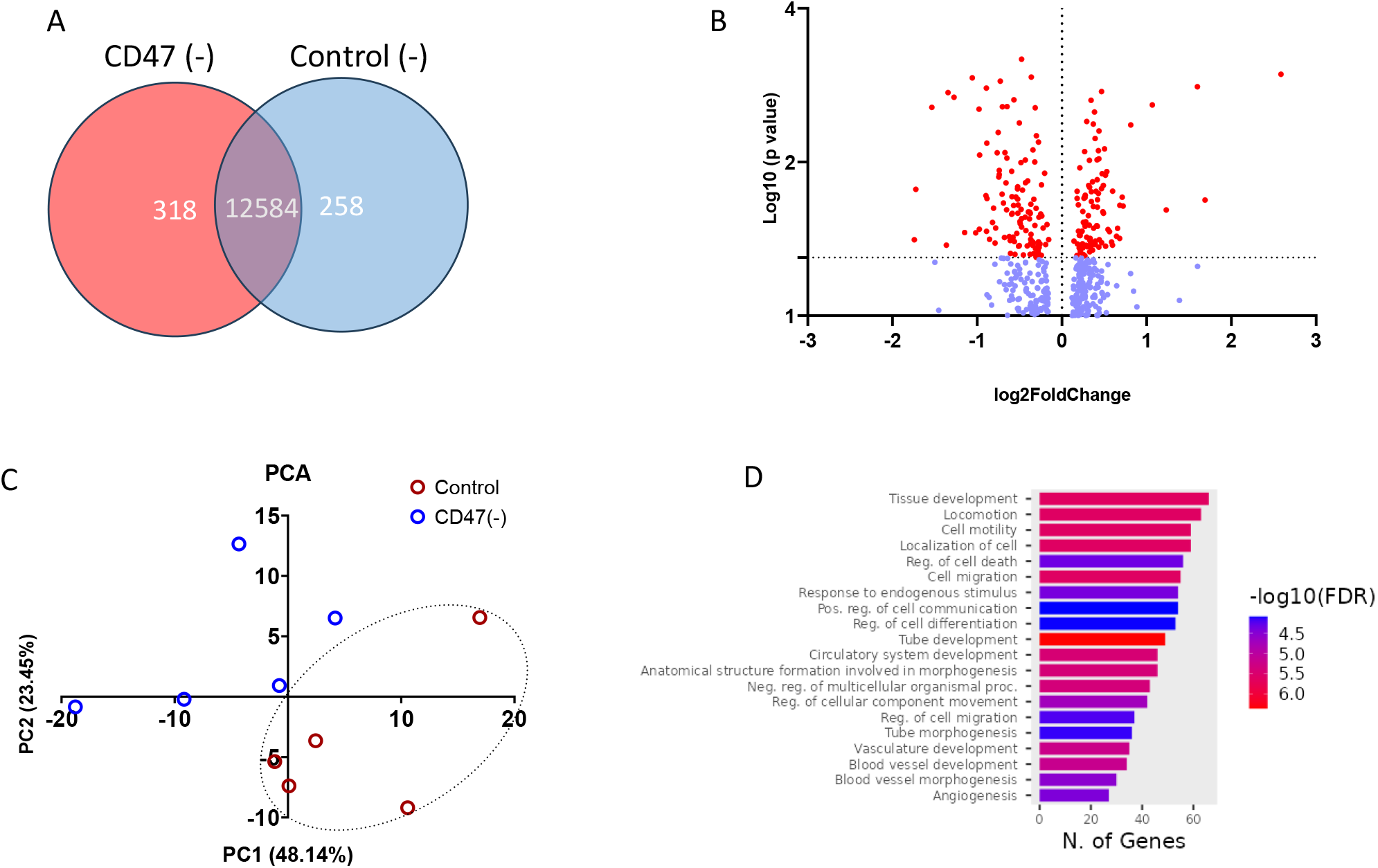
Blockade of CD47 regulates tumor gene expression. RNA sequencing was performed on 4T1-Br3 tumors of mice treated with isotype or anti-CD47 antibody. (A) Co-expression Venn Diagram represents the number of genes uniquely expressed within each group. (B) Volcano plot; horizontal axis for the fold change of genes in different samples. Vertical axis for a statistically significant degree of changes in gene expression levels. (C) Principal component analysis calculated by using discrete correlation summation (D) Significantly enriched terms in the GO enrichment analysis.

**Figure 3.**
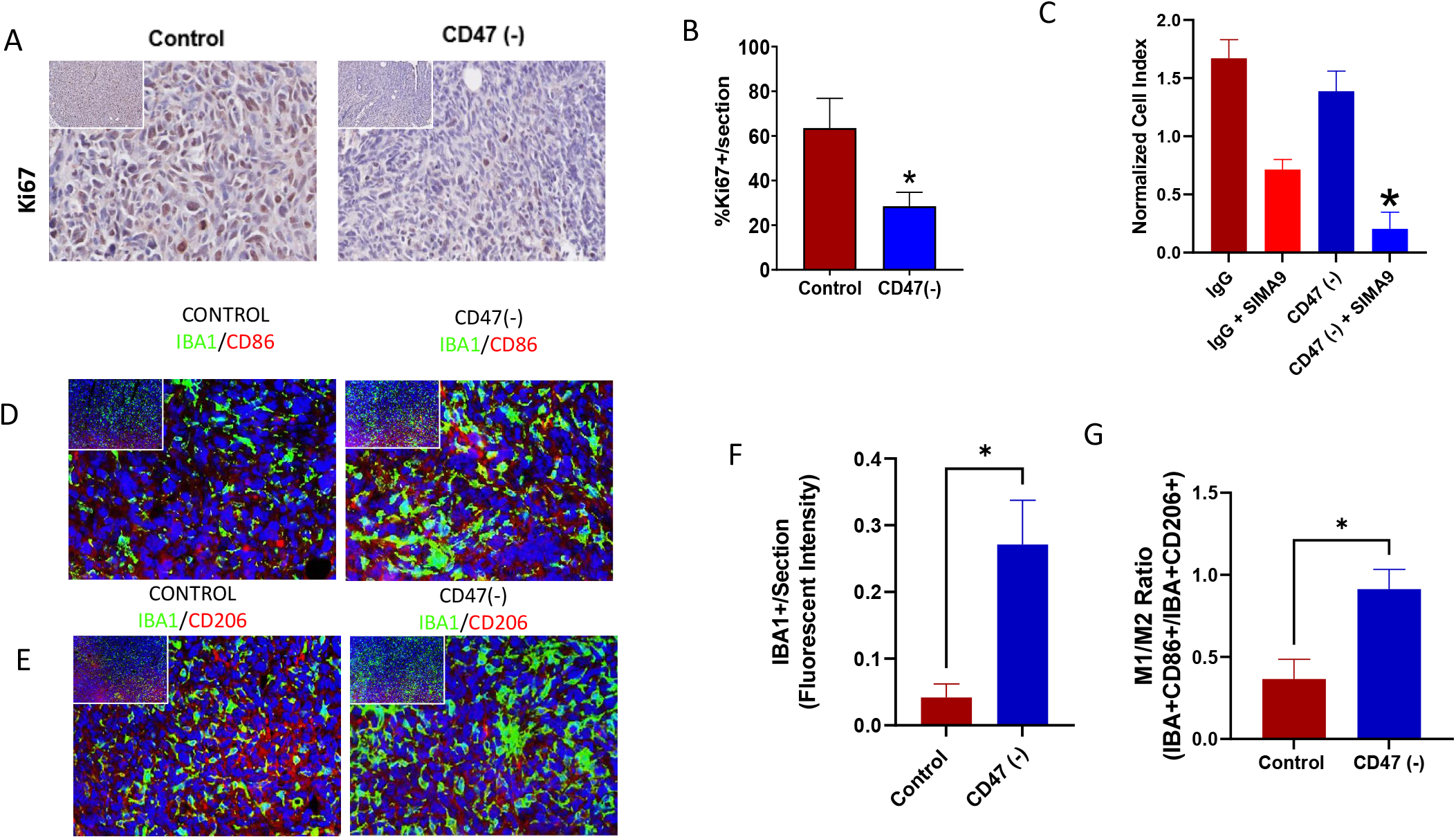
Effect of CD47 blockade on tumor proliferation and macrophage polarization. Tumor sections were paraffin-embedded and subjected to immunohistochemistry (A) Ki67 immunoreactivity to measure cell proliferation (brown stain) and (B) tissue segmentation analysis under BF analysis (n=6, *p<0.05). (C) Real-Time measurement of Microglia (SIM-A9) mediated cancer cell cytotoxicity (4T1-Br3) using cell impedance system (n=4). *p<0.05; **p<0.003. Immunofluorescence staining of in vivo orthotopic tumors with (D) IBA1-FITC/CD86-APC to detect M1-like macrophages or (E) IBA1-FITC/CD206-APC to detect M2-like macrophages. (F) Fluorescent intensity of IBA1 fluorescent staining alone (n=5, *p<0.05). (G) M1/M2 like macrophage ratios based on IBA1+ cells double positive for CD86 or CD206 (n=5, *p<0.05).

### Blockade of CD47 modulates cell viability in the tumor microenvironment

The top enriched pathway modulated by CD47 treatment includes pathways related to tissue development, including several pathways related to cell viability and proliferation; another significantly enriched pathway was cell death mechanisms. Therefore, to validate these observations, we stained tumor tissue sections with the proliferation marker Ki67 (Figure 3A). Our data show that blockade of CD47 reduced proliferating cells by 55.1% (63.5 ± 6.65 vs. 28.5 ± 3.12; % Ki67+ cells/section in IgG control vs. CD47(-) tumors; p=0.003; n=4) when compared to control group (Figures 3A & 3B). Another pathway by which CD47 is well known to induce cytotoxicity is through macrophage clearance. Since microglia are the main effector monocytes in the brain microenvironment, we co-cultured SIMA-A9 cells with 4T1-Br3 cells and measured microglial cell-mediated cancer cell cytotoxicity. Real-time measurement of microglia-mediated cytotoxicity revealed that CD47 blockade improves microglia-mediated cytolysis of 4T1-Br3 brain metastatic cancer cells by 68% (0.623 ± 0.07 vs. 0.202 ± 0.09 cell index of IgG+SIMA9 vs. CD47blockade+SIMA9; p=0.01; n=3, Figure 3 C). Since polarization of monocytes is key to understanding mechanisms of microglia-mediated cytotoxicity, we wanted to determine the relative amounts of M1 and M2 macrophages. Immunofluorescence staining for macrophage/microglial marker IBA1 revealed a 6.5-fold increase in macrophage infiltration in tumors treated with CD47(301) (1.00 ± 0.49 vs. 6.49 ± 1.59; Fold-change in IgG vs. CD47(301); p=0.017; n=4). Additionally, CD47(301) increased the macrophage M1/M2 ratio in tumors 2.5-fold (0.366 ± 0.12 vs. 0.914 ± 0.12; M1/M2 ratio in IgG vs. CD47(301); p=0.018; n=4). Thus, consistent with our RNA analysis, the reduction in tumor growth observed by CD47 blockade is associated with modulation of cell proliferation and cytotoxicity mediated by microglia.

### CD47 blockade limits brain trophic breast cell migration to reduce brain metastatic brain lesion formation

Our enrichment analysis demonstrated several pathways associated with breast cancer metastasis, including locomotion, cell motility, and cell migration. Therefore, we targeted CD47 on the 4T1b3 cell and plated the cells in a modified Boyden chamber well, and assessed migration by cell impedance (Figure 4A). We observed that targeting CD47 inhibits the migration of 4T1-Br3 brain trophic TNBC cells by 93% (11.59 ± 1.84 vs. 0.861 ± 0.25; normalized cell index of control vs. CD47 (-); p=0.001; n=4), Figures 4B and 4C). To determine whether CD47 blockade would regulate metastatic spread, we developed an in vivo model of breast-to-brain metastasis (Figure 4D). Female Balb/c mice were injected with 2 x10^5^ 4T1-Br3-Luc by intracardiac injection. Mice were treated with IgG or CD47 mAb every other day for one week. (Figure 4D). IVIS imaging on day 18 revealed that CD47 blockade reduced brain lesion formation by 88% (36775 ± 10366 vs. 4266 ± 1466; IVIS luminescence of IgG vs. CD47mAb; p=0.009; n=4-Figure 4E). Furthermore, we carried out a second tumor model using the brain trophic EO771-Br4 cell line to determine if the CD47 null microenvironment would impact metastatic spread. Thus, female C57BL/6 WT or CD47 null mice (-/-) were injected with 2.5 x 10^5^ EO771-Br4-Luc by intracardiac injection. (Add treatments-unsure of protocol) CD47 null mice had a significant improvement in overall survival (41%) compared to WT (0%) by day 35 (p=0.013). IVIS imaging revealed a % decrease in brain lesion formation in CD47 null mice (don’t have this graph – only picture). Additionally, there was a 2-fold increase in the microglia marker IBA1 in CD47 null anterior brain tissue (1.124 ± 0.29 vs. 2.429 ± 0.36; % IBA1+ cells/section in WT vs. CD47-/-; p=0.029; n=4). Thus, our data show that targeting CD47 o limiting its expression in the tumor microenvironment reduces metastasis of breast cancer cells to the brain by either autonomously reducing cancer cell migration or activating factors in the tumor microenvironment that clear or control migrating cancer cells.

**Figure 4.**
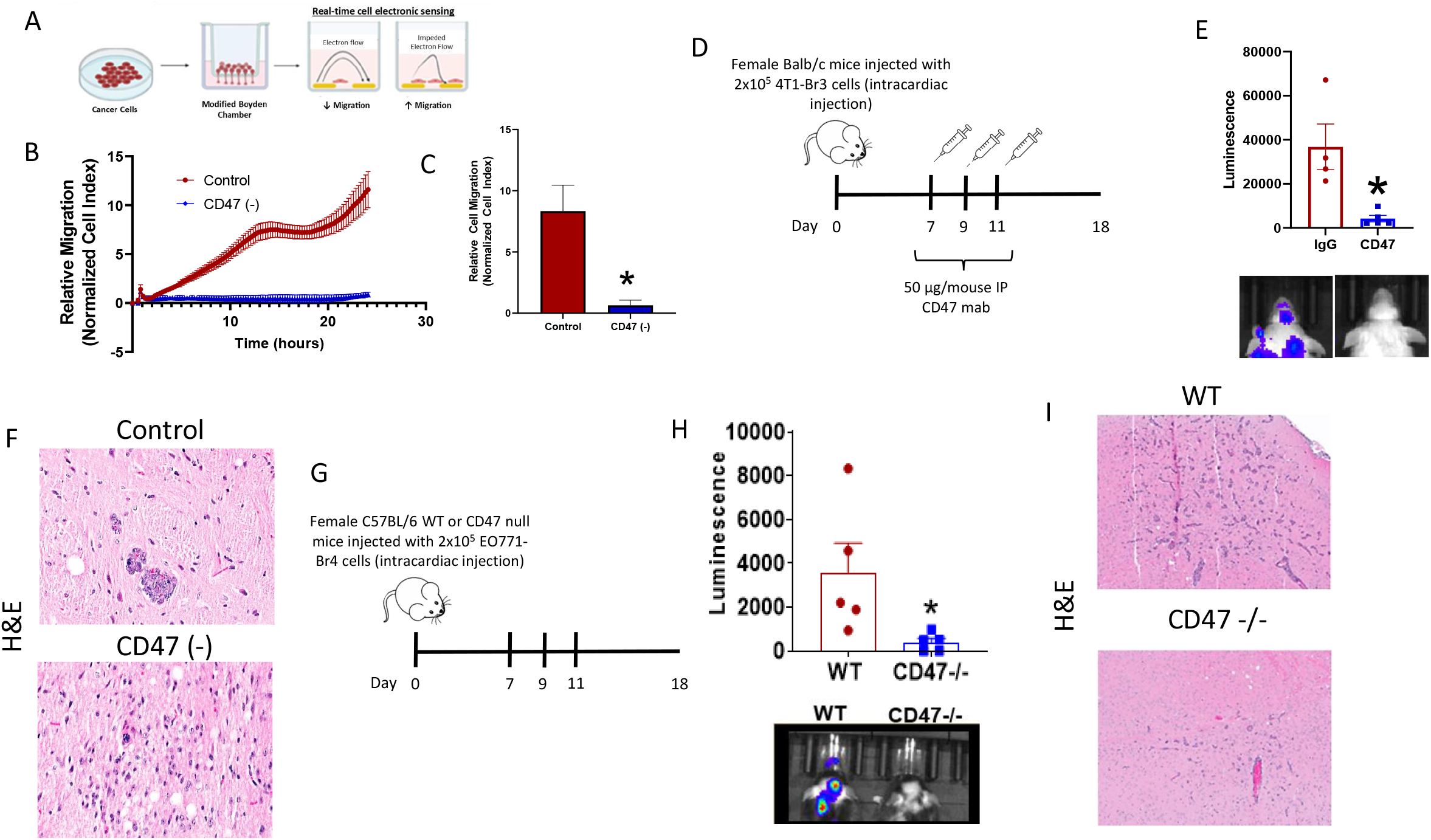
Blockade of CD47 inhibits migration and metastasis in vivo. (A) Schematic of cell migration measured using a modified Boyden chamber using the xCelligence Impedance system (scheme designed using biorender.com). (B) Representative plot of real-time relative migration calculated as normalized cell index and (C) quantification at 20h (n=3, *p<0.05). (D) Schematic of In vivo model of brain metastasis by intracardiac injection of 4T1-Br3-Luc cells; (E) quantification of luminescence intensity by IVIS imaging (n=4-5). (F) Hematoxylin and Eosin (H&E) staining of the anterior brain to confirm tumor lesions in control (IgG) or mice treated with anti-CD47 antibody (CD47(-)). (G) schematic of EO771-Br4 intracardiac model in WT or CD47 null (CD47-/-) mice. (H) quantification of luminescence intensity by IVIS imaging (n=5, *p<0.05). (I) confirmation of tumor lesions by H&E staining.

## Discussion

One of the major complications of breast cancer is the metastatic spread to brain. TNBC is associated with a higher risk of metastasis, and the brain is one of the common sites of distant metastases in this subtype. The clinical management of breast-to-brain metastasis includes surgery, radiation therapy, and systemic therapies. Still, many patients succumb to disease, which has led to the incorporation of immunotherapy approaches in managing brain metastatic breast cancer. The combination of chemotherapy and anti-PD-1 is approved for treating invasive breast cancer; however, whether this approach is feasible remains under investigation (24). Therefore, novel immunotherapy targets are needed to improve patient outcomes.

CD47 is an innate and adaptive immune checkpoint. While the role of CD47 in cancer is widely explored in cancer, its role in the progression from breast to brain metastasis is not well understood. We and others have demonstrated that CD47 is overexpressed in invasive breast cancer (25,26) and that targeting CD47 reduces TNBC tumor and metastasis to the lung in a preclinical syngeneic model(23). In this report, we show that expression of CD47 is elevated in metastatic lesions from patients with metastatic breast cancer compared to normal tissue. Furthermore, CD47 expression is significantly elevated in the bran trophic 4T1 cell line when compared to the parental, suggesting that this receptor would be a potential target for breast cancer brain metastasis.

The role of CD47 in metastasis is mainly attributed to immune escape limiting innate anti-tumor immunity (27). In the brain tumor microenvironment, CD47 restricts the effector action of microglia against glioblastoma lesions (28,29), and signaling is known to be implicated in monocyte polarization (30). While we understand that the signaling of glioblastoma and metastatic brain lesions differs, these studies suggest potential implications of CD47 signaling in metastatic brain cancer progression. Indeed, our data show that targeting CD47 increases the M1/M2 microglia ratio in our in vivo models and that targeting this receptor enhances microglia-mediated cytotoxicity of brain trophic breast cancer cells. While this data is not surprising, the role of CD47 in metastatic escape, particularly to the brain, is limited and thus critical to develop strategies for this complication in breast cancer patients.

Aside from mechanisms of innate anti-tumor immunity, our studies demonstrated that blockade of CD47 reduced brain trophic breast cancer cell migration. This is consistent with studies implicating that increased expression of CD47 is associated with the spread and invasiveness of several cancer types (27). Previous works indicate that MAPK pathway activation is associated with the CD47 effect on epithelial cell migration and that inhibition of Src reverses this effect (31). Our RNAseq data of genes downregulated by anti-CD47 treatment indicated significant enrichment for cell migration and motility genes. These included the downregulation of Matrix Metalloproteinases that have been well-documented to be involved in cancer progression and metastasis (32). A study examining neurovascular injury associated with stroke showed that the interaction of TSP1 and CD47 induces activation of MMP-9 in brain endothelial cells and astrocytes. Furthermore, treatment with the TAX2 peptide, which is reported to inhibit the interaction of TSP1 and CD47, limits melanoma metastasis, and other studies using this peptide have demonstrated inhibition of MMP signaling (33,34). Thus, our observations of MMP downregulation are likely related to antibody treatment or the absence of CD47 in the microenvironment limiting activation by TSP1.

One key aspect that our data revealed by our model in CD47 null mice is that the absence of CD47 in the tumor microenvironment also reduces metastatic burden in the brain. Most studies on CD47 signaling focus on its expression on cancer cells and the bypassing of anti-tumor immunity or ligand/receptor interactions mediated intracellular signaling of cancer cells. However, our model using EO771 brain trophic cells, which are WT for CD47, supports the role of CD47 affecting non-cancer cells to potentially promote tumor progression. We have previously observed that targeting CD47 on T cells alone enhanced effector function against fibrosarcoma cells. Implantation of B16 melanoma tumors in CD47 null mice also decreased tumorigenesis(17), which was associated with an increase in granzyme B. Furthermore, later studies demonstrated that TSP1 limited T cell activation through CD47 by specifically limiting metabolic reprogramming needed to promote cytotoxic T cell effector function (18). T cell populations are limited in the brain microenvironment. Still, in the context of metastasis and disruption of the blood-brain barrier the population of lymphocytes can increase, which is one of the potential reasons that immune checkpoint inhibitors are relatively more effective in the brain metastatic setting when compared to glioblastoma (35). Since microglia are the main effector cells in the brain microenvironment, the blockade of CD47 could also modulate the metabolic reprogramming of these cells to enhance the clearance of metastatic lesions.

Immune-targeting approaches have limited activity in invasive breast cancer. Our data show that CD47 may be a driver of metastatic progression and that targeting these receptors significantly reduces metastatic burden in preclinical models of TNBC brain metastasis. We know that a single agent is unlikely to show this type of activity in human patients; still, since antibodies to CD47 are in clinical trials for other malignancies, this study could motivate trials using combinatorial strategies, including standard of care whole brain irradiation to treat brain metastatic breast cancer. The radiation approach in combination with CD47 blockaded would be supported by our previous studies and later studies by others demonstrating synergism with radiotherapy(16,36). Overall, we are confident in the strong anti-tumor activity observed in our study, and thus incorporating CD47 blockade in the treatment strategy of metastatic brain cancer could result in the significant enhancement of survival of breast cancer patients.

## References

1. Perou CM, Sorlie T, Eisen MB, van de Rijn M, Jeffrey SS, Rees CA, et al. Molecular portraits of human breast tumours. Nature 2000;406:747–52

2. Lv Y, Ma X, Du Y, Feng J. Understanding Patterns of Brain Metastasis in Triple-Negative Breast Cancer and Exploring Potential Therapeutic Targets. Onco Targets Ther 2021;14:589–607

3. Al-Mahmood S, Sapiezynski J, Garbuzenko OB, Minko T. Metastatic and triple-negative breast cancer: challenges and treatment options. Drug Deliv Transl Res 2018;8:1483–507

4. Brosnan EM, Anders CK. Understanding patterns of brain metastasis in breast cancer and designing rational therapeutic strategies. Ann Transl Med 2018;6:163

5. Howlader N, Cronin KA, Kurian AW, Andridge R. Differences in Breast Cancer Survival by Molecular Subtypes in the United States. Cancer Epidemiol Biomarkers Prev 2018;27:619–26

6. Ribas A, Wolchok JD. Cancer immunotherapy using checkpoint blockade. Science 2018;359:1350–5

7. Sharma P, Allison JP. Immune checkpoint targeting in cancer therapy: toward combination strategies with curative potential. Cell 2015;161:205–14

8. Jahchan NS, Mujal AM, Pollack JL, Binnewies M, Sriram V, Reyno L, et al. Tuning the Tumor Myeloid Microenvironment to Fight Cancer. Front Immunol 2019;10:1611

9. Liu X, Kwon H, Li Z, Fu Y-X. Is CD47 an innate immune checkpoint for tumor evasion? Journal of Hematology & Oncology 2017;10

10. Liu X, Pu Y, Cron K, Deng L, Kline J, Frazier WA, et al. CD47 blockade triggers T cell-mediated destruction of immunogenic tumors. Nat Med 2015;21:1209–15

11. Kauder SE, Kuo TC, Harrabi O, Chen A, Sangalang E, Doyle L, et al. ALX148 blocks CD47 and enhances innate and adaptive antitumor immunity with a favorable safety profile. PLoS One 2018;13:e0201832

12. Advani R, Flinn I, Popplewell L, Forero A, Bartlett NL, Ghosh N, et al. CD47 Blockade by Hu5F9-G4 and Rituximab in Non-Hodgkin’s Lymphoma. N Engl J Med 2018;379:1711–21

13. Weiskopf K. Cancer immunotherapy targeting the CD47/SIRPalpha axis. Eur J Cancer 2017;76:100–9

14. Chung H, Lee K, Kim W, Gainor J, Lakhani N, Chow L, et al. ASPEN-01: A phase 1 study of ALX148, a CD47 blocker, in combination with trastuzumab, ramucirumab and paclitaxel in patients with second-line HER2-positive advanced gastric or gastroesophageal junction cancer. Ann Oncol 2021;32:S215–S6

15. Kim TM, Lakhani N, Gainor J, Kamdar M, Fanning P, Squifflet P, et al. A Phase 1 Study of ALX148, a CD47 Blocker, in Combination with Rituximab in Patients with Non-Hodgkin Lymphoma. Blood 2019;134

16. Maxhimer JB, Soto-Pantoja DR, Ridnour LA, Shih HB, DeGraff WG, Tsokos M, et al. Radioprotection in normal tissue and delayed tumor growth by blockade of CD47 signaling. Sci Transl Med 2009;1:3ra7

17. Soto-Pantoja DR, Terabe M, Ghosh A, Ridnour LA, DeGraff WG, Wink DA, et al. CD47 in the tumor microenvironment limits cooperation between antitumor T-cell immunity and radiotherapy. Cancer Res 2014;74:6771–83

18. Stirling ER, Terabe M, Wilson AS, Kooshki M, Yamaleyeva LM, Alexander-Miller MA, et al. Targeting the CD47/thrombospondin-1 signaling axis regulates immune cell bioenergetics in the tumor microenvironment to potentiate antitumor immune response. J Immunother Cancer 2022;10

19. UniProt C. UniProt: the universal protein knowledgebase in 2021. Nucleic Acids Res 2021;49:D480–D9

20. Wixon J, Kell D. The Kyoto encyclopedia of genes and genomes--KEGG. Yeast 2000;17:48–55

21. Ge SX, Jung D, Yao R. ShinyGO: a graphical gene-set enrichment tool for animals and plants. Bioinformatics 2020;36:2628–9

22. Bronson SM, Westwood B, Cook KL, Emenaker NJ, Chappell MC, Roberts DD, et al. Discrete Correlation Summation Clustering Reveals Differential Regulation of Liver Metabolism by Thrombospondin-1 in Low-Fat and High-Fat Diet-Fed Mice. Metabolites 2022;12

23. Feliz-Mosquea YR, Christensen AA, Wilson AS, Westwood B, Varagic J, Melendez GC, et al. Combination of anthracyclines and anti-CD47 therapy inhibit invasive breast cancer growth while preventing cardiac toxicity by regulation of autophagy. Breast Cancer Res Treat 2018;172:69–82

24. Nakhjavani M, Shigdar S. Natural Blockers of PD-1/PD-L1 Interaction for the Immunotherapy of Triple-Negative Breast Cancer-Brain Metastasis. Cancers (Basel) 2022;14

25. Kaur S, Elkahloun AG, Singh SP, Arakelyan A, Roberts DD. A function-blocking CD47 antibody modulates extracellular vesicle-mediated intercellular signaling between breast carcinoma cells and endothelial cells. J Cell Commun Signal 2018;12:157–70

26. Yan H, Huang W, Chen C, Zhang X, Zhu K, Yuan J. MiR-133a/CD47 axis is a novel prognostic biomarker to promote triple negative breast cancer progression. Pathol Res Pract 2023;244:154400

27. Lian S, Xie X, Lu Y, Jia L. Checkpoint CD47 Function On Tumor Metastasis And Immune Therapy. Onco Targets Ther 2019;12:9105–14

28. Hutter G, Theruvath J, Graef CM, Zhang M, Schoen MK, Manz EM, et al. Microglia are effector cells of CD47-SIRPalpha antiphagocytic axis disruption against glioblastoma. Proc Natl Acad Sci U S A 2019;116:997–1006

29. Cole AP, Hoffmeyer E, Chetty SL, Cruz-Cruz J, Hamrick F, Youssef O, et al. Microglia in the Brain Tumor Microenvironment. Adv Exp Med Biol 2020;1273:197–208

30. Zhang M, Hutter G, Kahn SA, Azad TD, Gholamin S, Xu CY, et al. Anti-CD47 Treatment Stimulates Phagocytosis of Glioblastoma by M1 and M2 Polarized Macrophages and Promotes M1 Polarized Macrophages In Vivo. PLoS One 2016;11:e0153550

31. Shinohara M, Ohyama N, Murata Y, Okazawa H, Ohnishi H, Ishikawa O, et al. CD47 regulation of epithelial cell spreading and migration, and its signal transduction. Cancer Sci 2006;97:889–95

32. Roy R, Morad G, Jedinak A, Moses MA. Metalloproteinases and their roles in human cancer. Anat Rec (Hoboken) 2020;303:1557–72

33. Jeanne A, Boulagnon-Rombi C, Devy J, Theret L, Fichel C, Bouland N, et al. Matricellular TSP-1 as a target of interest for impeding melanoma spreading: towards a therapeutic use for TAX2 peptide. Clin Exp Metastasis 2016;33:637–49

34. Li H, Xu H, Wen H, Wang H, Zhao R, Sun Y, et al. Lysyl hydroxylase 1 (LH1) deficiency promotes angiotensin II (Ang II)-induced dissecting abdominal aortic aneurysm. Theranostics 2021;11:9587–604

35. Di Giacomo AM, Mair MJ, Ceccarelli M, Anichini A, Ibrahim R, Weller M, et al. Immunotherapy for brain metastases and primary brain tumors. Eur J Cancer 2023;179:113–20

36. Soto-Pantoja DR, Sipes JM, Ghosh A, Merino MJ, Roberts DD. Therapeutic targeting of CD47 regulates cell bioenergetics and autophagy to reduce breast tumor growth and protect against anthracycline-mediated cardiac toxicity. 2014; San Diego, CA. Cancer Res.

